# Real-time Visualization of Trigger Factor on Translating Ribosomes

**DOI:** 10.1101/2025.05.16.654432

**Authors:** Eider Nuñez, Prithwidip Saha, Markel G.Ibarluzea, Arantza Muguruza-Montero, Sara M-Alicante, Rafael Ramis, Aritz Leonardo, Aitor Bergara, Alvaro Villarroel, Felix Rico

## Abstract

Trigger Factor (TF) is an ATP-independent chaperone that assists in co-translational protein folding by associating with ribosomes to prevent aggregation. While its interaction with ribosomes has been described, real-time visualization of TF dynamics has remained elusive. Using high-speed atomic force microscopy, we imaged full 70S ribosomes under near-physiological conditions during translation. TF exhibited dynamic transitions between extended and compact conformations, forming both stable and transient contacts with ribosomal proteins uL23 and bL17 in ribosome–nascent chain complexes. Binding to non-translating ribosomes was not observed under these conditions. Molecular dynamics simulations of TF alone and in complex with ribosomal proteins supported the experimental observations. Our findings reveal the structural flexibility of TF and its selective association with active ribosomes. Our combination of experimental and computational approaches offers new insights into how TF dynamically engages ribosomes during translation to facilitate protein folding.

## Introduction

Protein synthesis is a fundamental cellular process in which ribosomes translate messenger RNA (mRNA) into polypeptides that must then fold into functional conformations. Ribosomes, complex macromolecular machines made of ribosomal RNA (rRNA) and proteins, are the sites of protein synthesis. In prokaryotes, they consist of two subunits: the large (50S) and the small (30S) subunit. These subunits work together to bind messenger RNA (mRNA) and transfer RNA (tRNA) during translation. As the ribosome reads the mRNA, it synthesizes the corresponding polypeptide, which begins to fold within the ribosomal exit tunnel^1^.

Preventing protein misfolding or aggregation is imperative for maintaining cellular homeostasis and ensuring proper protein function. Chaperones play a pivotal role in facilitating co-translational folding, ensuring proteostasis by assisting nascent polypeptides in achieving their correct structures and preventing aggregation-prone intermediates^2-4^.

Trigger factor (TF) is an essential chaperone in bacteria. It associates with the ribosome at the exit tunnel and interacts with nascent polypeptide chains as they emerge from the ribosome^5-10^. The interaction with ribosomal proteins uL23 and bL29, located adjacent to the exit tunnel, is vital in preventing premature folding or aggregation of nascent chains, ensuring their proper folding only after translation is completed^11-13^.

Beyond its role in nascent chain folding, TF also acts as a protective factor under stress conditions, stabilizing the ribosome and ensuring the fidelity of protein synthesis^10,14^. TF is composed of three primary domains: the N-terminal ribosome binding domain (RBD), the peptidyl-prolyl cis/trans isomerase (PPIase) domain, and the C-terminal substrate-binding domain (SBD)^11^. The RBD and SBD are integral to its chaperone function, facilitating the stabilization and proper folding of nascent polypeptides as they exit the ribosome^15,16^. While the PPIase domain contributes to its overall structure, it is not essential for its chaperone activity *in vivo*^17-19^.

Cryo-electron microscopy (cryo-EM) and X-ray crystallography have provided valuable static structures of TF bound to the ribosome^10,11^. However, these techniques are limited in capturing the dynamic nature of interactions occurring during translation. Recent advancements in single-molecule techniques and computational methods have shown that TF exhibits significant conformational flexibility, adapting to the nascent chain and the ribosome’s functional state^20^. However, the lack of real-time observations under physiological conditions limits our understanding of how these interactions occur in situ. Specifically, it remains unclear how TF dynamically engages with different ribosomal partners, how its conformational changes relate to function, and how selectivity for translating ribosomes is achieved.

Advances in high-speed atomic force microscopy (HS-AFM) have enabled real-time visualization of macromolecular complexes in solution with nanometer resolution^21-25^. HS-AFM overcomes the limitations of complex sample preparation and artificial stabilization techniques, providing a more accurate representation of macromolecular complex dynamics. The study by Imai et al. (2020)^25^ represents a pivotal step in advancing our understanding of ribosomal dynamics. Their work reported real-time visualization of GTPase binding to the P-stalk of the large ribosomal subunit, offering new insights into ribosome-GTPase dynamics and showcasing HS-AFM’s potential for studying transient molecular events. However, the use of only the 50S subunit with a specially engineered chimera containing the P-stalk from Archaea may not fully reflect physiological conditions due to structural modifications. Additionally, their analysis focused solely on the 50S subunit, limiting observations of the complete ribosome during protein synthesis and its interaction with other proteins like TF.

Here, we use HS-AFM to directly visualize TF dynamics and interactions with translating ribosomes under near-physiological conditions. We captured TF’s conformational transitions in real-time and observed stable and transient interactions with ribosomal proteins uL23 and the less-characterized bL17. Our data uncover novel aspects of TF engagement and conformational plasticity. Complementary molecular dynamics (MD) simulations further support our findings, providing atomic-resolution insights. By visualizing both the localization and dynamic behavior of TF in situ, without the need for engineered constructs, our study establishes a new standard for probing ribosome-associated chaperones in native contexts.

## Results

### 1. High-Speed AFM visualization of intact ribosomes in solution

To preserve ribosomal integrity during HS-AFM imaging, we optimized buffer composition, surface preparation, and imaging parameters to enable full ribosome visualization. We isolated native 70S ribosomes from *Escherichia coli* using an optimized two-step protocol, comprising a sucrose cushion followed by a sucrose gradient, designed to preserve their structural integrity and functionality (see Materials & Methods). The MgAc-containing imaging buffer stabilized the ribosomes without causing excessive subunit dissociation, ensuring a balanced environment that maintained their native conformations during imaging^26-29^. Bare mica proved ideal for partially immobilizing ribosomes, facilitating effective electrostatic interactions between the negatively charged surface and MgAc-stabilized ribosomes. Under these conditions, we successfully visualized intact ribosomes under near-physiological conditions, capturing their structural dynamics in various orientations, as shown in Fig. 1 and Supplemental Videos 1 to 5.

**Figure 1.**
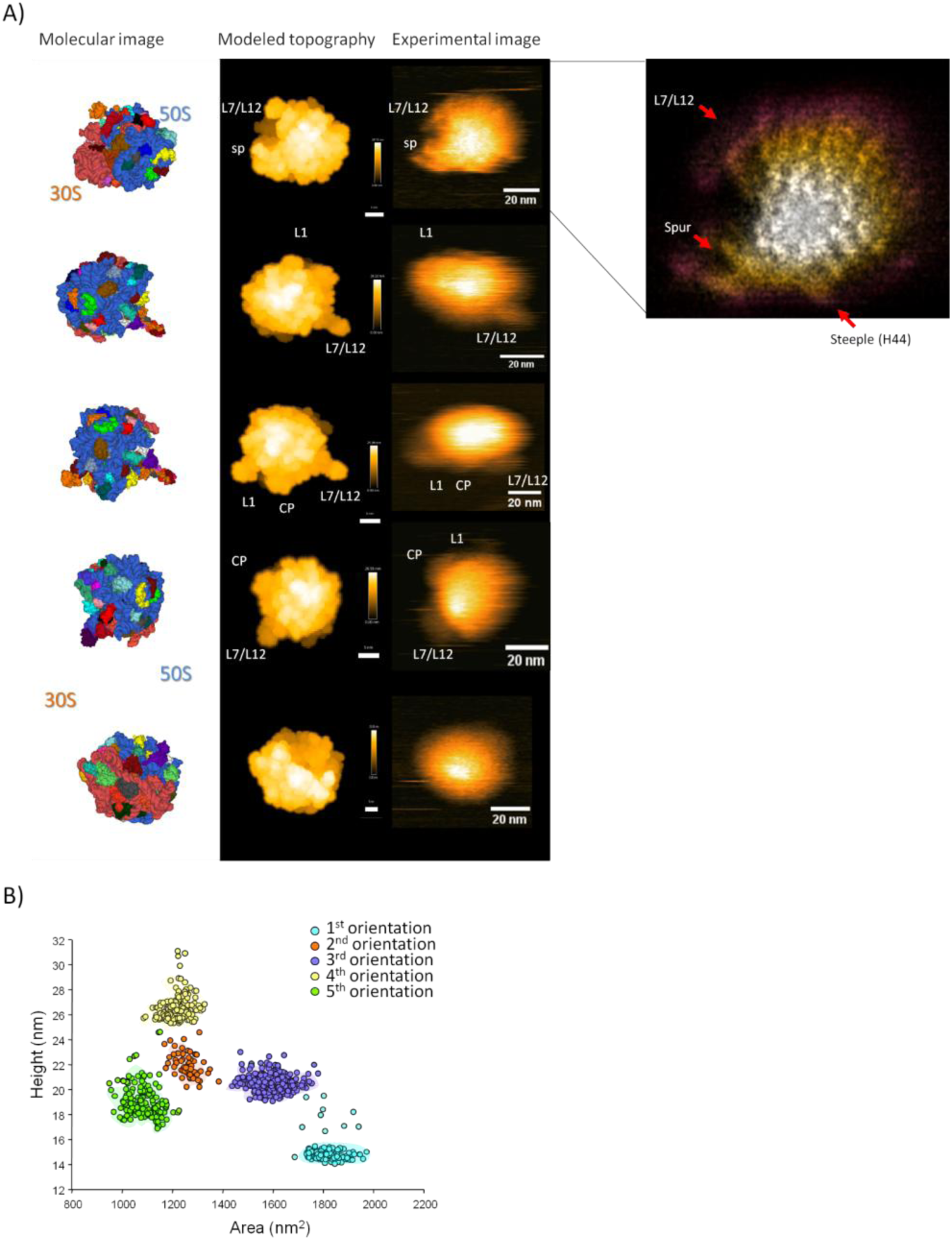
Visualization of the 70S Ribosome in Four Different Orientations on a Mica Surface. **(A)** Experimental HS-AFM images of the 70S ribosome in various orientations. Molecular images and BioAFMviewer-modeled topographies were generated from the structure of the translating bacterial ribosome (PDB: 7K00)^33^. In the molecular image, the 50S subunit is highlighted in blue, while the 30S subunit appears in coral. Key structural features are labeled: sp = spur, L1 stalk, CP = central protuberance, and L7/L12 = P-stalk base. On the right, a map of the first orientation generated using a localized AFM algorithm applied over multiple frames. **(B)** Graphical representation of the ribosome’s height and diameter measurements across different orientations shown in panel A.

#### Structural Insights into Ribosome Dynamics

Using BioAFMviewer^30,31^, we modeled atomic force microscopy (AFM) topography images from the bacterial ribosome structure at 2.0 Å resolution (PDB:7K00)^32^, predicting five distinct ribosomal orientations based on the most probable interactions with the mica surface. Since the L7/L12 stalk was omitted in this PDB structure, we incorporated it into the model. This adjustment allowed for a more accurate comparison between modeled and experimental images (Fig. 1, Supplemental Videos 1–5).

Distinct orientations revealed key ribosomal features, including the spur of the 30S subunit and the L7/L12 stalk and central protuberance (CP) of the 50S subunit (Fig. 1A). Height and projected area measurements provided further insights into ribosomal conformational states (Fig. 1B). The 70S ribosome exhibited an average height of 20.5 ± 3.3 nm, while the isolated 50S subunit measured 15.2 ± 1.2 nm (Supplemental Fig. 1A), consistent with previous reports of hydrated ribosomes^25,33^.

In the first orientation (Panel A, Fig. 1), the ribosome appeared in a lying position with a height of 15.3 nm, corresponding to the maximal height of the 50S subunit (Supplemental Fig. 1A). This view emphasized the head of the 30S subunit and revealed structural variability even in regions lacking direct functional roles, such as the spur, which corresponds to helix 6 of the 16S rRNA (Supplemental Fig. 1B). Interestingly, coarse-grained simulations by Liu and Zhang (2023)^34^ suggest that this observed mobility could hold functional significance, possibly influencing ribosomal dynamics during translation.

To enhance resolution, we applied the localization AFM algorithm over multiple frames to generate an LAFM image^35,36^ (Fig. 1A, right). Although precise identification of specific features remains challenging when compared with Cryo-EM data, this approach allowed us to distinguish protein and RNA domains within the ribosome. In the LAFM reconstruction, we observed a protrusion that may correspond to the *steeple* of h44 (Fig. 1A, right), a feature that remained consistently visible across multiple datasets. Further comparisons with Cryo-EM structures of *E. coli* ribosomes could help clarify its identity and potential role in this context.

The second and third orientation clearly distinguished the CP and the P-stalk, with the latter again displaying significant flexibility (Supplemental Fig. 1B), consistent with previous observations of ribosomal subunit dynamics^25,37,38^(Fig. 1, Supplemental Videos 2–3). The fourth orientation also revealed stalk base movement (Fig. 1, Supplemental Video 4), aligning with Imai et al. (2020)^25^. However, while Imai et al. reported a broader motion range (80° to 240°) in a chimeric ribosome combining the *E. coli* 50S subunit with the archaeal P-stalk^25^, our measurements in the unmodified *E. coli* ribosome revealed a more restricted angular transition of 90° to 130° (Supplemental Fig. 1C). This reduced mobility may be attributed to the presence of the 30S subunit in the complete ribosome of our data, as opposed to the isolated large subunit analyzed by Imai et al. The additional steric and dynamic constraints imposed by interactions within the intact ribosome could limit the conformational freedom of the P-stalk. Differences in experimental conditions, buffer composition, or the chimeric nature of the P-stalk in the Imai et al. study could also contribute to the observed discrepancies. In this orientation, the L1 stalk is also distinguishable in some frames (Supplemental Video 4).

The fifth ribosomal orientation showed the small subunit positioned over the large subunit, giving the ribosome a spherical-like view, where individual features became harder to distinguish (Fig. 1, Supplemental Video 5). Interestingly, we also observed ribosomal disassembly. As the small subunit detached, only the large subunit remained visible, underscoring the transient nature of ribosomal interactions (Supplemental video 5 and Supplementary Fig. 1E).

The average area was 2010 ± 720 nm² for the 70S ribosome and 1040 ± 340 nm² for the 50S subunit (Fig. 1B, Supplemental Fig. 1D), supporting the distinction between complete and partial ribosomal forms under HS-AFM. While tip convolution effects inherent to AFM did introduce some measurement distortions, they did not obscure clear differentiation between ribosomal forms. Histograms illustrating height and area distributions captured structural states and orientations, providing real-time visualization of ribosomal architecture (Supplemental Fig. 1A).

### 2. Characterization of TF monomers using HS-AFM

With the final goal of exploring the interaction of the TF with ribosomes, we first characterized the conformational dynamics of isolated TF monomers in solution using HS-AFM imaging at a concentration of 5 nM, well below the dimerization dissociation constant (KD range of 1–18 µM)^39-41^. While resolved structures of the TF monomer have been derived from the dimeric form or from TF bound to the ribosome^14,42-45^, the monomeric form of TF in solution has not been resolved. Our HS-AFM approach allowed us to observe real-time structural transitions of the TF monomer, providing valuable novel insights into its behavior in solution.

#### Structural Dynamics and Flexibility of TF Monomer

The average area of the monomer was 110 ± 17 nm², and its height was 6 ± 1 nm (Supplemental Fig. 2A), consistent with the expected dimensions for the monomeric form from the crystal structure, which was reported as three domains folded in an elongated conformation^42^. Capturing one frame per second, we observed significant conformational flexibility over an 80-second observation period (Fig. 2A). To quantify this conformational flexibility, we calculated the aspect ratio of the particles, revealing continuous fluctuations, from 1.1 to 2.0. Extended, anisotropic conformations were characterized by higher ratios (1.55–2.00), with approximately 27.8% of frames falling within this range. The majority of frames (37.7%) displayed more compact conformations with ratios between 1.00 and 1.19, while 34.5% showed intermediate values (1.20–1.54) (Fig. 2B, Supplemental Video 6). The shifts in aspect ratio observed highlight the inherent flexibility of the TF monomer as it undergoes conformational cycling between extended and compact states, a key feature of its functional mechanism^10,16,46^.

**Figure 2.**
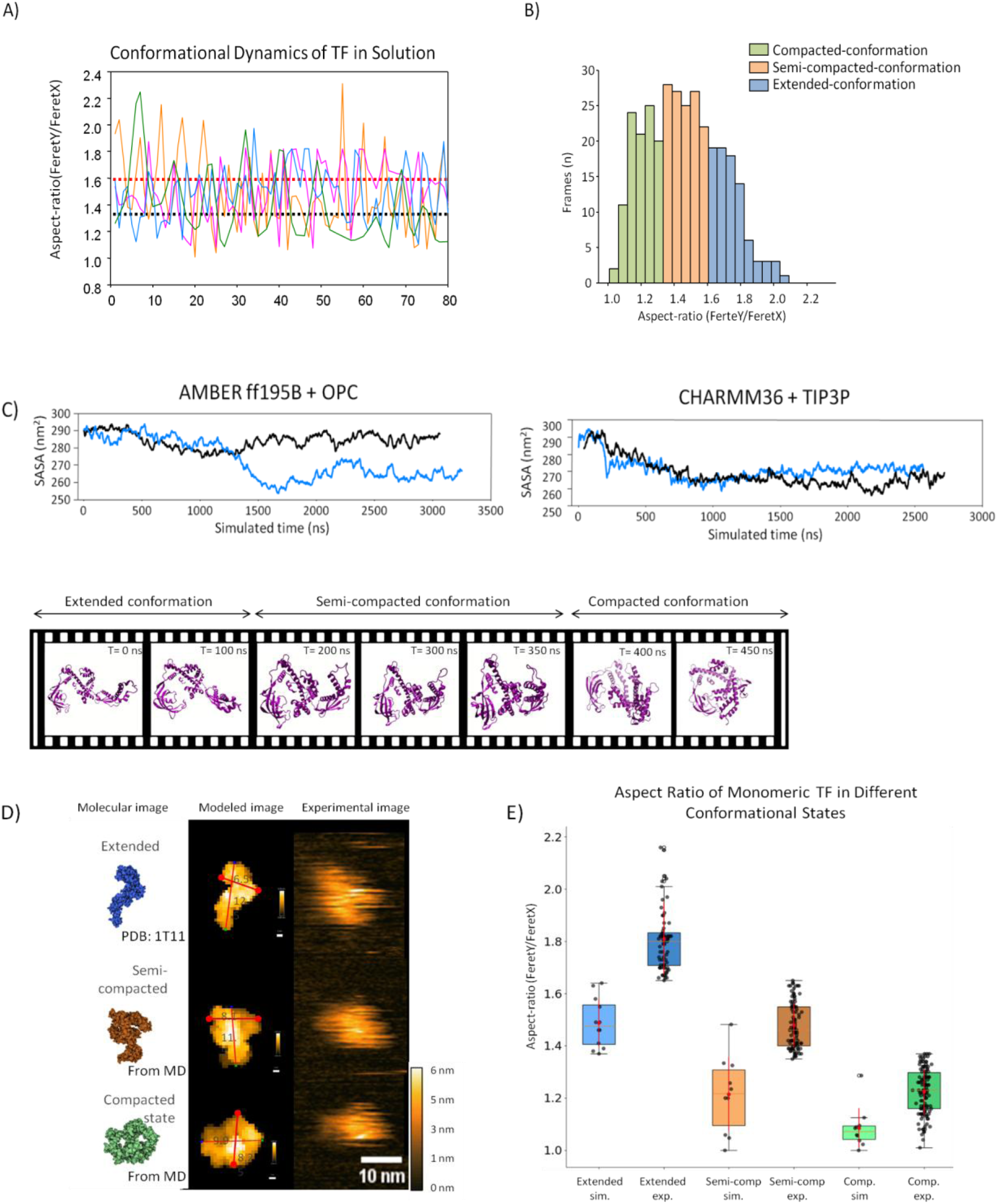
Conformational Flexibility of the TF Monomer in Solution. **(A)** Time-dependent variation in the aspect ratio of the TF monomer at a concentration of 5 nM in physiological buffer (25 mM Tris, pH 7.5, 120 mM KCl, 5 mM NaCl, 14 mM MgAc). Data from different experiments are represented in distinct colors. **(B)** Histogram of conformational state distributions. **(C)** Top: Solvent Accessible Surface Area (SASA) plots of the TF monomer over time from four independent simulations. The left panel presents results obtained using the Amber ff19SB + OPC force field, while the right panel shows results from the CHARMM36 + TIP3P force field. Bottom: frames from MD simulations showing the compaction trajectory of TF. **(D)** Molecular representations of three distinct TF conformations: extended (top, initial state, PDB: 1T11), semi-compacted (middle, at 1400 ns of CHARMM36 simulation), and collapsed (bottom, CHARMM36 simulation endpoint). Simulated topographies were generated with BioAFMviewer. **(E)** Boxplots of simulation and experimental aspect ratio measurements for the extended (blue), semi-compacted (orange), and compacted (green) conformations. Simulation aspect ratios are represented in solid colors, while experimental data are shown in lighter shades, with individual data points displayed for each state. Simulation values were calculated in BioAFMviewer and analyzed across at least 10 different orientations. Experimental aspect ratios were determined by dividing FeretY by FeretX, with classification thresholds of >1.6 for the extended state, 1.65–1.35 for the semi-compacted state, and <1.37 for the compacted state.

#### Molecular Dynamics Simulations of TF Compaction

To investigate the mechanisms driving the conformational compaction of TF, we conducted MD simulations using the Amber ff19SB and CHARMM36 protein force fields, paired with the OPC and TIP3P water models, respectively, revealing similar dynamic behaviors. The simulations were based on the monomeric TF structure derived from PDB entry 1T11^42^. Each simulation spanned approximately 3 µs, (Fig. 2C, Supplemental videos 7 and 8). We monitored the solvent-accessible surface area (SASA), which reflects the extent of surface exposed to the solvent. The temporal dynamics of SASA revealed a compaction process. A decrease in SASA was observed as TF transitioned from an extended to a more compact state, indicating an increased burial of hydrophobic regions and greater stability of the tertiary structure (Fig. 2C). The sequential frames in Fig. 2C (bottom) visually depicts this structural transition.

#### Distinct Conformational States in MD Simulations

Three distinct conformational states of TF—extended (E), semi-compacted (SC), and compacted (C)— were identified, each exhibiting unique structural and dynamic properties. Representative conformations of these states were extracted from MD simulations and compared with experimental data (Fig. 2D). The extended conformation was modeled based on the PDB structure 1T11, while the semi-compacted and compacted states emerged from the MD simulations at 1400 ns and at the simulation endpoint, respectively. Topographic models of these conformations, generated using BioAFMviewer, were juxtaposed with experimental HS-AFM images for structural validation. Notably, the compacted state had not been previously characterized in experimental studies, nor is there a resolved PDB structure for this conformation. Our MD simulations revealed this conformation, which exhibited a reduced aspect ratio and closely matched the real-time structural transitions observed via HS-AFM (Fig. 2D).

To quantitatively assess these structural states, simulation aspect ratios were computed using BioAFMviewer across at least 10 molecular orientations, considering electrostatic interactions with the mica surface, charge distribution, and molecular topology. The calculated aspect ratios were 1.6 ± 0.1 for the extended state, 1.2 ± 0.2 for the semi-compacted state, and 1.1 ± 0.1 for the compacted state (Fig. 2E). Experimental aspect ratios followed the classification thresholds of >1.6 for the extended state, 1.65–1.35 for the semi-compacted state, and <1.37 for the compacted state, yielding mean values of 1.8 ± 0.2, 1.5 ± 0.1, and 1.2 ± 0.1, respectively. Simulation values were in excellent agreement with experimental measurements (Fig. 2E), further reinforcing the structural correspondence between simulations and experimental observations. This agreement provides strong computational-experimental validation for the existence of a compacted TF state, reinforcing the structural correspondence between the two approaches.

#### Comparison of TF Monomer and Dimer by HS-AFM

To promote dimer formation, the concentration of TF was increased to 5 µM (Supplemental video 9). The dimer exhibited a larger molecular area (241 ± 89 nm²) and height (8 ± 1 nm) compared to the monomer (101 ± 17 nm², 6 ± 1 nm) (Supplemental Fig. 2A). This substantial increase in area and height reflects the structural rearrangements and added volume associated with dimer formation. The observed variability in the dimer area suggests a dynamic equilibrium, where not only dimers but also higher-order oligomers, such as trimers or tetramers, may form, as previously described for TF from *Thermotoga maritima*^47^ (Supplemental Fig. 2A). The aspect ratio of the dimer remained relatively stable over time, ranging from 1.3 to 1.5 (Supplemental Fig. 2B), while the monomer exhibited a broader range, fluctuating from 1.1 to 2.2 (Fig. 2A). These observations suggest that the dimer adopts a more stable structure with reduced conformational flexibility. This finding is consistent with previous studies, which proposed that the TF dimer exists in a conformationally dynamic equilibrium while maintaining overall structural stability^44^.

### 3. Interaction of Trigger Factor with Ribosomes

We then imaged the dynamic binding of TF to ribosomes, both in the presence and absence of a nascent polypeptide chain. In the absence of a nascent chain TF exhibited minimal ribosome binding, even at a 1000:1 TF-to-ribosome ratio (Supplemental Fig. 3 A), Supplemental Video 10). To obtain a homogeneous population of translating ribosomes, we introduced a SecM arrest peptide at the C-terminus of monomeric Teal Fluorescent Protein 1 (mTFP1), which stalled translation at a specific point (Fig. 3A). This positioning placed mTFP1 at the N-terminus of the arrested protein, outside the ribosomal exit tunnel, forming a distinct protuberance in ribosome images and enabling us to distinguish translating from non-translating ribosomes.

**Figure 3.**
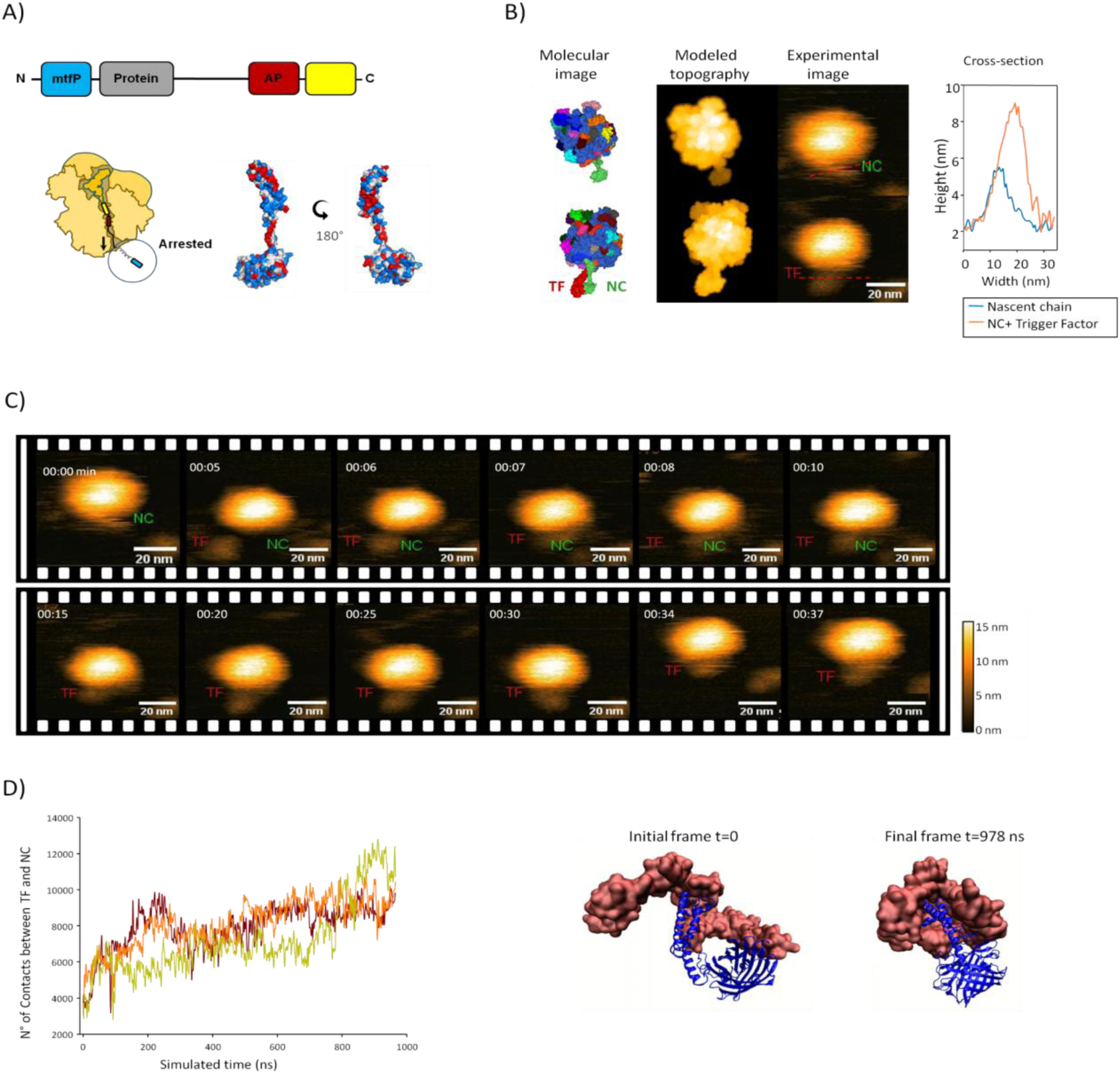
Trigger Factor (TF) Binding to the Ribosome Nascent Chain Complex (RNC). **(A)** Schematic representation of the nascent chain used in the TF binding assay. The construct includes a blue mTFP1 fluorescent protein, gray α-helices (A, TW, and B) from the KCNQ channel, a red SecM (Ec-Ms) arrest peptide, and a yellow mcpVenus protein (not translated). The AlphaFold2 structure of the accessible portion of the nascent chain is shown, colored by hydrophobicity (red for hydrophobic, blue for hydrophilic, and white for neutral residues). **(B)** Diagram of the TF-RNC interaction, displaying molecular images of the ribosome complex with and without TF binding. Modeled AFM topographies from BioAFMViewer compared to experimental data, alongside a graph showing cross-sectional topography for the nascent chain alone versus TF-bound. **(C)** Sequential frames illustrating TF binding to RNC in a physiological buffer (RNC at 2 nM, TF at 1 μM), showing that TF remains bound for over30 seconds. **(D)** Molecular dynamics simulations depicting the TF-NC interaction. The number of contacts between the nascent chain and TF is tracked over 1000 ns using a cutoff of 3 Å. On the right, the initial and final frames of the interaction between the RNC (blue) and TF (surface, red) reveal how TF embraces the nascent chain.

To further characterize this system, we employed AlphaFold2^48^ to model the structure of mTFP1 outside the ribosomal exit tunnel (Fig. 3A). The model revealed specific hydrophobic patches within its α-helices, which have been proposed as key TF binding sites^49,50^. This experimental setup provided a biologically relevant framework for investigating TF binding dynamics under conditions that closely resemble the *in vivo* environment.

The measured height of the ribosome-nascent chain (RNC) particles was approximately 17 ± 3 nm, consistent with the dimensions of the large ribosomal subunit. The nascent chain exhibited a cross-sectional height of ∼5 nm, in agreement with estimates for the nascent chain or the AlphaFold model, and revealed important flexibility (Fig. 3B).

TF binding to the nascent chain was evidenced by an increase in cross-sectional height to 9.6 nm, suggesting that TF stabilizes the structure of the emerging polypeptide. This height increase is consistent with TF-polypeptide association. Additionally, the lateral dimensions of the complex expanded, with the width increasing from 11 to 20 nm, further supporting TF binding (Fig. 3B, Supplemental Video 11). Monomeric binding supports the flexible engagement mechanism observed in our molecular dynamics simulations, highlighting TF’s ability to adapt its conformation to the nascent chain.

TF binding to the RNC complex persisted for more than 30 seconds, as shown in Fig. 3C and Supplementary Video 11. To quantify the interaction kinetics, we modeled the dwell time of TF on the ribosome using a simple exponential distribution. The fit yielded a characteristic time constant (τ) of ∼8 seconds (Fig. 3C and Supplementary Fig. S3). This value closely matched the trend observed in the histogram of dwell times. Notably, our experimental setup involved reconstituted systems with ribosomes stalled on mRNAs, a methodology that may not fully capture the dynamic interaction of TF with the growing nascent chain in the cellular context.

To investigate the molecular determinants of TF binding to the nascent polypeptide, we conducted all-atom MD simulations over 1000 ns in three independent replicates (Fig. 3D, Supplementary Video 12). Throughout the simulations, TF established and maintained extensive interactions with the nascent chain, stabilizing more than 10,000 atomic contacts per replicate by the end of the trajectory. These interactions were predominantly localized to the hydrophobic regions of the nascent chain, aligning with TF’s well-established role in shielding aggregation-prone sequences during translation (Fig. 3D). As the simulation progressed, TF gradually enveloped the nascent polypeptide, suggesting a dynamic recognition mechanism in which TF adapts its conformation to stabilize emerging hydrophobic motifs (Fig. 3D, initial and final frames). This adaptive engagement is consistent with previous biochemical and structural studies, as well as our own HS-AFM observations of TF flexibility, demonstrating TF’s flexible substrate recognition and stabilization^51^.

The C-terminal domain of TF (residues 248–432) was critical in mediating these interactions, indicating a sustained hydrophobic binding interaction. This finding aligns with prior structural and biochemical evidence showing that TF specifically recognizes and binds exposed hydrophobic patches on nascent polypeptides to prevent misfolding^14,52^. To further characterize these interactions, we quantified the minimum distance between the C-terminal domain of TF and the nascent chain, generating a histogram that revealed a predominant minimum distance of 2.6 Å, with values ranging from 2.45 to 3.55 Å (Supplementary Fig. 4A). Additionally, the simulations indicated an increase in hydrogen bonding interactions as TF engaged with the nascent chain, further stabilizing the emerging structure and enhancing the recognition process (Supplementary Fig. 4B). This increase in hydrogen bonds likely contributes to the stabilization and proper folding of the nascent polypeptide.

### 4. Stable and transient binding patterns of the TF to RNC

We identified two sequential binding sites for TF on RNC complexes occurring simultaneously, suggesting that the binding stoichiometry may not always be 1:1, as previously thought^6,7^. The first, stable binding mode, lasting over 30 seconds, was characterized by the RBD of TF anchoring to uL23 on the large ribosomal subunit, followed by its embrace of the nascent chain (Fig. 4C, Supplemental Video 11). Subsequently, we observed a second and transient interaction near ribosomal protein bL17, which lasted less than 15 seconds (Fig. 4B and 4C, Supplementary Fig. 5 and Supplemental Videos 11, 13, 14). These interactions were detected in RNCs visualized from different orientations (Fig. 4A). To characterize their kinetics, we modeled the dwell times using exponential distributions. The first TF binding event yielded the mentioned time constant of τ = 8.0 ± 1.1 seconds, while the second, more transient interaction near bL17 showed a significantly shorter time constant of τ = 3.9 ± 0.4 seconds (Supplementary Fig. 3B). This finding is significant because it provides the first direct visualization of transient interactions with bL17, previously hypothesized by Martinez-Hackert and Hendrickson (2009)^43^, and now experimentally captured in our study. These results support a model where TF engages ribosomal proteins in a sequential and modular manner, possibly tuning its activity according to the stage of nascent chain emergence.

**Figure 4.**
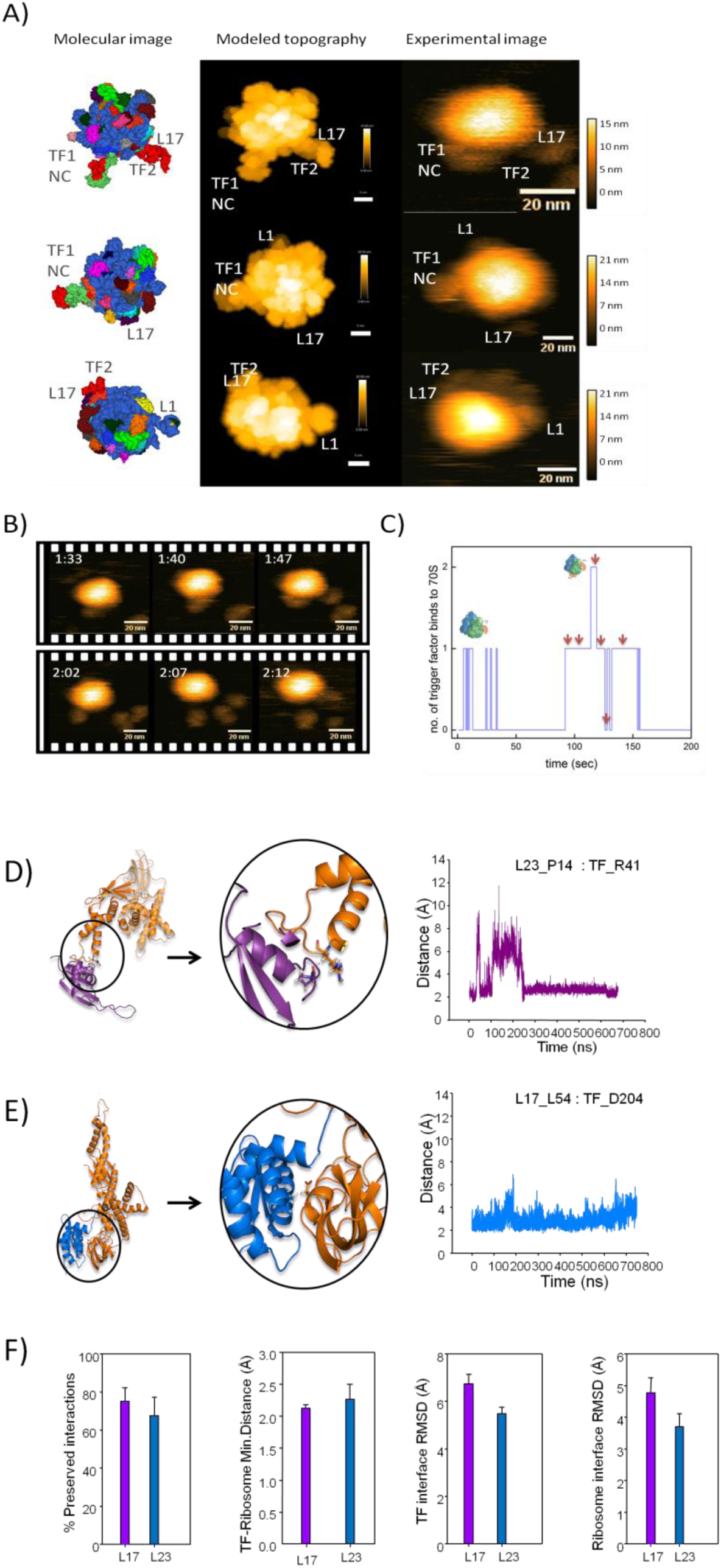
Simultaneous Binding of Multiple TF Molecules to the Ribosomes. **(A)** HS-AFM image alongside molecular and simulated topography. AFM-simulated images are based on the 50S ribosomal subunit (PDB: 6PJ6), with the nascent chain (NC) predicted by AlphaFold2 (green) and Trigger Factor (TF) (red). **(B)** Sequential frames showing two TF molecules dynamically binding to the ribosome. **(C)** Graph depicting the number of TF molecules bound over time in (B), illustrating the frequency and duration of binding events. Red arrows indicate the frames shown in (B). **(D)** Interaction of ribosomal proteins uL23 and bL17 with TF, analyzed through molecular dynamics simulations. Left: Schematic representation of the interaction between L23 and TF, highlighting L23 P14 and TF R41. Right: Graph showing the distance between these residues over ∼700 ns of simulation. **(E)** Schematic representation of the interaction between L17 and TF, emphasizing L17 D54 and TF D204. The accompanying graph shows the distance between these residues over ∼800 ns of simulation. **(F)** Graphs summarizing the percentage of preserved L17-TF and L23-TF interactions, distances between these protein pairs, RMSD of the TF interface, and RMSD of the ribosomal protein interface across all analyzed systems.

#### Molecular Dynamics Simulations: Binding Specificity and Structural Adaptability

To analyze the interactions between TF and the ribosomal proteins bL17 and uL23 at atomic resolution, we performed MD simulations spanning ∼800 ns. The simulations revealed that TF maintained stable interatomic distances below 5 Å with both bL17 and uL23, indicating persistent physical contact (Fig. 4D and E; Supplemental Videos 15 and 16). The distance plots in these panels show contacts between TF R41 and uL23 P14 (Fig. 4D), and between TF D204 and bL17 D54 (Fig. 4E), both of which were preserved over hundreds of nanoseconds, lending support to the structural feasibility of these interactions.

Residue-level analysis identified key contact points. The PPIase domain of TF (residues D204–G208) predominantly engaged with bL17 (residues L54–A61) through hydrophobic stacking and hydrogen bonding (Fig. 4E). Although this interaction does not involve the catalytic site, it may serve as a non-catalytic anchoring mechanism that positions TF near the ribosomal exit tunnel. Conversely, the RBD of TF (residues R41–D43) formed stable contacts with uL23 (residues P14–V16), supporting a role in initial tethering of TF to the ribosome and facilitating its early chaperone function^41,47^(Fig. 4D).

Root mean square deviation (RMSD) analysis of the TF residues at the interfaces revealed distinct interaction dynamics. The TF–uL23 interface exhibited lower RMSD values (3.7–4.0 Å), consistent with a stable interaction, whereas the TF–bL17 interface showed higher flexibility (6.7–7.0 Å). This contrast suggests that the TF–uL23 interaction may represent an initial anchoring point, while TF–bL17 may act as a transient scanning or repositioning contact. Such functional asymmetry aligns with the proposed sequential binding model and may enable TF to adjust to the conformational state of the ribosome and nascent chain.

Interaction persistence was quantified at 75% for bL17 and 67% for uL23 (Fig. 4F). Interactions were defined by a minimum interatomic distance ≤6 Å. While these values suggest stable interactions, we note that persistence percentages alone may overestimate functional significance; thus, complementary validation using site-directed mutagenesis or crosslinking assays would be beneficial in confirming the biological relevance of these interfaces.

Additionally, RMSD of the ribosome upon TF binding remained low (∼5.5 Å for bL17 and ∼5.8 Å for uL23), suggesting that TF does not induce major conformational rearrangements in the ribosomal structure. However, this metric may underestimate localized dynamic effects, particularly at the exit tunnel, which could influence co-translational folding events or the recruitment of other factors. Overall, these findings support a model in which TF acts as a conformationally flexible chaperone, capable of dynamic repositioning on the ribosome to engage emerging nascent chains without compromising ribosomal integrity^44^.

## Discussion

Using HS-AFM, we captured dynamic conformational states and interactions of bacterial ribosomes and the TF chaperone under near-physiological conditions. Our images confirm the inherent flexibility of ribosomal substructures—particularly the L1 stalk, P-stalk, and 30S spur—supporting previous hypotheses^53^.

We observed that the TF monomer dynamically transitions among extended, semi-compact, and compact states. MD simulations corroborated these findings, revealing a preference for compact conformations. These transitions occur on physiologically relevant timescales and align with the chaperone cycle described previously^54-56^. While earlier studies focused on dimeric or ribosome-bound TF, our data reveal structural features of the free monomer, validating extended states and uncovering semi-compact and compact forms that had only been hypothesized previously^56^. These results, together with simulations and RNC interaction data, highlight the TF monomer’s conformational plasticity as essential for interacting with diverse substrates, independently of ATP hydrolysis.

The semi-compact states may represent intermediate conformations relevant for folding, as proposed by Thomas et al. (2013)^46^, potentially stabilizing partially folded proteins or mediating transient contacts. This supports the idea that the monomer’s intrinsic flexibility enables its role as a versatile chaperone. In contrast, the dimeric form shows limited flexibility, with a stable aspect ratio (1.3–1.5) and higher area and height, in agreement with previous reports^44^. This rigidity may serve functional roles, preserving a defined cavity for substrate binding and preventing nonspecific interactions.

Previous studies have documented stable interactions between TF and non-translating ribosomes^58,59^. However, we did not observe stable TF interactions with non-translating ribosomes, but instead found consistent binding to translating ribosomes, reinforcing the model that TF recognition is driven by emerging hydrophobic segments of nascent chains. These findings align with earlier data indicating substrate-dependent TF affinity and are further supported by live-cell tracking studies showing that TF binding to non-translating ribosomes is transient, falling below the detection limit of millisecond time resolution^10,47,52,60^.

To probe TF-RNC interactions, we used an engineered nascent chain fused to mTFP1 and the SecM arrest peptide. High-resolution models identified hydrophobic α-helical patches as TF binding sites, consistent with TF’s known preference for exposed hydrophobic regions^50,61^.

MD simulations revealed that TF forms stable, long-lasting interactions with the nascent polypeptide over a 1000 ns simulation period. The C-terminal domain of TF plays a key role in recognizing and stabilizing hydrophobic segments within the nascent chain, underscoring its function in shielding aggregation-prone regions^16,62,63^. The close contact between TF and specific nascent chain residues highlights its dual role as both a chaperone and a scaffold during co-translational folding^64^.

We observed a stable interaction between the nascent chain and arm 2 of the C-domain of TF (Supplemental Fig. 4C), consistent with previous studies that identified this region as a primary binding site for nascent chains^65^, reinforcing its central role in their recognition and stabilization.

The stable and transient interactions of TF with ribosomes, observed through MD simulations and HS-AFM, provide valuable insights into its binding dynamics. Specifically, the stable interaction between the TF-RBD and uL23, persisting for over 30 seconds. In contrast, the interaction of TF with ribosomal protein bL17 was markedly more transient, with a characteristic dwell time of only 3.9 ± 0.4 seconds (Supplementary Fig. 3B). Both interactions were specific to translating ribosomes. These observations suggest TF uses a dual mechanism: stable anchoring via uL23 and transient scanning via bL17, possibly for regulating folding or quality control. This finding is consistent with the work of Yang et al. (2022)^66^, who identified bL17 as part of a hydrophobic ribosomal patch that interfaces with emerging polypeptides. Given TF’s affinity for hydrophobic sequences, its association with bL17 may reflect a dynamic docking mechanism that modulates co-translational folding or processing. Although the precise functional significance of these brief encounters remains to be fully elucidated, their transient nature could allow TF to scan the ribosomal surface efficiently or to engage selectively with specific nascent chain sequences.

RMSD measurements of TF interactions with ribosomal proteins bL17 and uL23 provide deeper insight into the adaptability of TF’s binding dynamics. The TF-uL23 interface exhibits greater stability (3.7–4.0 Å), in contrast to the more flexible TF-bL17 interaction (6.7–7.0 Å), highlighting a functional balance between structural rigidity and dynamic flexibility. Such a balance allows TF to anchor nascent chains effectively while maintaining the flexibility needed for transient interactions during co-translational folding, without disrupting translation. Notably, minimal changes in ribosomal conformation (∼5.5 Å at bL17 and ∼5.8 Å at uL23) upon TF binding further support its role as a peripheral chaperone, aiding protein maturation without interfering with ribosomal function^44^. Furthermore, the interaction between the TF-PPIase domain (D204-G208) and bL17 (L54-A61) has now been directly observed, providing the direct evidence for this hypothesized contact, previously suggested by biochemical studies^43^.

Emerging evidence indicates that TF binding to RNCs is more transient in vivo than in vitro. Our experimental observations, along with those from other in vitro studies using stalled ribosomes, likely reflect the absence of continuous nascent chain elongation and ribosomal translocation—factors that promote TF dissociation in the cellular context. These differences underscore the need for further research combining both in vitro and in vivo approaches to fully elucidate how TF coordinates co-translational folding within the dynamic and crowded cellular environment. To address this, future studies should aim to develop more physiologically relevant in vitro systems that mimic ribosomal elongation and translocation for observation in real time^60,66^. Our study places real-time observation of protein synthesis within reach.

## Materials and Methods

### 1. Sample expression and purification

#### a. 70 S ribosome purification from *E.coli* bacteria

To purify 70S ribosomes from *Escherichia coli*, cells were cultured in LB medium (Thermo Fisher Scientific, #12780052) at 37°C until reaching an OD600 of 0.5–0.7, corresponding to mid-logarithmic growth. Cells were harvested by centrifugation at 5,000 × g for 10 minutes at 4°C to maintain ribosome integrity. The resulting pellets were washed with ice-cold PBS (Gibco, #70011036) and stored on ice until lysis.

Bacterial pellets were resuspended in lysis buffer (20 mM Tris-HCl, pH 7.5, 10 mM MgCl₂, 100 mM NH₄Cl, 0.5 mM EDTA, and 6 mM β-mercaptoethanol; Sigma-Aldrich, #M6250) and lysed using a high-pressure homogenizer (Avestin, EmulsiFlex-C5). The lysate was clarified by ultracentrifugation at 30,000 × g for 30 minutes at 4°C to remove unlysed cells and large debris.

The resulting supernatant was subjected to a high-salt wash with buffer containing 20 mM Tris-HCl (pH 7.5), 500 mM NH₄Cl, 10 mM MgCl₂, and 6 mM β-mercaptoethanol to eliminate loosely associated proteins. The washed supernatant was layered onto a 30% sucrose cushion prepared in ribosome stabilization buffer (20 mM Tris-HCl, pH 7.5, 10 mM MgCl₂, 100 mM NH₄Cl, and 6 mM β-mercaptoethanol) and centrifuged at 100,000 × g for 16–18 hours at 4°C using a Beckman Coulter ultracentrifuge with a SW32 Ti rotor.

The ribosome pellet was carefully resuspended in low-salt buffer (20 mM Tris-HCl, pH 7.5, 10 mM MgCl₂, 50 mM NH₄Cl, and 6 mM β-mercaptoethanol) to avoid aggregation. For further purification, the resuspended pellet was layered onto a 10–40% sucrose gradient in the ribosome stabilization buffer and ultracentrifuged at 150,000 × g for 5 hours at 4°C.

Ribosomal fractions were collected by monitoring absorbance at 260 nm to identify the 70S ribosome peak. Collected fractions were analyzed by SDS-PAGE (NuPAGE™ 4-12% Bis-Tris Protein Gels, Invitrogen, #NP0321BOX) to confirm purity. Purified ribosomes were aliquoted and stored at -80°C.

#### b. Trigger Factor purification

Trigger Factor (TF) protein was purified from *Escherichia coli* strain BL21 (DE3) cells (Novagen, #69450) transformed with a plasmid encoding His-tagged TF. Cultures were grown in LB medium (Thermo Fisher Scientific, #12780052) supplemented with 50 µg/mL kanamycin (Sigma-Aldrich, #K0254) at 37°C to an OD600 of 0.6–0.8. Protein expression was induced by adding 0.5 mM IPTG (Thermo Fisher Scientific, #15529019), and cultures were incubated for an additional 4 hours at 25°C to optimize protein folding.

Cells were harvested by centrifugation at 5,000 × g for 15 minutes at 4°C, and the pellet was washed with ice-cold PBS (Gibco, #70011036). The bacterial pellet was resuspended in lysis buffer (50 mM Tris-HCl, pH 7.5, 300 mM NaCl, 10 mM imidazole, and 1 mM PMSF) and lysed by sonication on ice with a Qsonica sonicator (model Q500) using 10 cycles of 10 seconds on, 20 seconds off. The lysate was clarified by centrifugation at 30,000 × g for 30 minutes at 4°C.

The supernatant was loaded onto a Ni²⁺-NTA affinity column (HisTrap HP, GE Healthcare, #17-5248-01) pre-equilibrated with a lysis buffer. Unbound proteins were removed by washing the column with a wash buffer (50 mM Tris-HCl, pH 7.5, 300 mM NaCl, 20 mM imidazole) until baseline absorbance returned to zero. TFr was then eluted with an elution buffer containing 50 mM Tris-HCl, pH 7.5, 300 mM NaCl, and 250 mM imidazole (Sigma-Aldrich, #I5513).

Eluted fractions were dialyzed overnight at 4°C against buffer (20 mM Tris-HCl, pH 7.5, 150 mM NaCl, 1 mM DTT) using a dialysis membrane with a 10 kDa cutoff (Thermo Fisher Scientific, #68100). Following dialysis, the sample was concentrated with Amicon Ultra-15 Centrifugal Filters (Millipore, #UFC901008) and further purified by size-exclusion chromatography on a Superdex 200 column (GE Healthcare, #28990944) equilibrated with a buffer (20 mM Tris-HCl, pH 7.5, 150 mM NaCl, 1 mM DTT).

Eluted fractions containing pure TF were confirmed by SDS-PAGE using NuPAGE™ 4-12% Bis-Tris gels (Invitrogen, #NP0321BOX) and stained with Coomassie Brilliant Blue (Bio-Rad, #1610436). The purified TF fractions were pooled, concentrated, and stored at -80°C in aliquots for subsequent analysis.

### 2. HS-AFM imaging

#### a. Sample preparation

A freshly cleaved muscovite mica disc (diameter: 1.5 mm; thickness: 0.1 mm; JBG-Metafix, Montdidier, France) was used as a surface without any further modification. The mica was then glued to a glass stage for imaging purposes. Ribosomes were prepared in an imaging buffer consisting of 120 mM potassium acetate (KAc), 25 mM Tris (pH 7.5), and 1 mM β-mercaptoethanol (β-Me), supplemented with 14 mM magnesium acetate (MgAc). A 2 µL aliquot of protein solution, at a concentration of 5 nM, was deposited onto the mica surface and incubated for 2-3 minutes to allow for partial immobilization of the ribosomes. Following incubation, the surface was washed several times with the imaging buffer to remove any unbound ribosomes. HS-AFM imaging was conducted in the same buffer environment, with 100 µL of buffer in the imaging chamber. For experiments involving TFs, it was introduced directly to the imaging chamber at a final concentration of 5 nM, followed by an incubation period of 10 minutes before imaging.

#### b. Data acquisition

Imaging was conducted in tapping mode on an SS-NEX HS-AFM equipped with a standard scanner and ultrashort cantilevers having a resonance frequency of 600 kHz in liquid and a nominal spring constant of 0.15 N/m (USC-F1.2-k0.15, NanoWorld, Neuch.tel, Switzerland). During scanning, the free oscillation amplitude of the cantilever was set to 3 - 4 nm and the set point for feedback control was kept 20 % lower than this value. Images were acquired at a scan rate of 1 frame per second. Typically, the pixel size was 0.5 ’ 0.5 nm^2^, while the typical size of a scan area was 300 ’ 300 nm^2^. At least 3 independent experiments were performed both for ribosome alone and ribosome in the presence of TF. All measurements were performed at room temperature (22 - 25 °C) in the imaging buffer.

#### c. Image processing and analysis

All HS-AFM videos were processed using laboratory-built macros implemented in Fiji (ImageJ) image processing software to semi-automate the image processing. The processing steps mainly involve noise reduction, background correction, and the removal of unwanted background particles. To detect the binding of the TF to the ribosome, we employed another macro to identify the bound TF as a protrusion on the ribosome surface across hundreds of images. The code used for image processing is available on the GitHub repository^67^. LAFM maps were carried out after alignment of the imaging frames using Template Matching and applying the localization algorithm provided in the original work on ImageJ^35,68,69^.

The aspect ratio was calculated using ImageJ by dividing the Feret X (the maximum width of the particle) by the Feret Y (the maximum height of the particle).

### 3. Molecular Dynamics Simulation

#### a. Trigger Factor Monomer Simulations

Molecular Dynamics (MD) simulations of the TF monomer (PDB 1W26) were performed using two force fields: CHARMM36^70^ with TIP3P water^71^ and Amber ff19SB^72^ with OPC water^73^. The CHARMM36 simulations were conducted using GROMACS 2020^74^, while Amber simulations were carried out using the pmemd program in Amber2270^75^.

The TF monomer was placed in a cubic simulation box with periodic boundary conditions and solvated in a cubic box with an ionic concentration of 0.15 M NaCl. For the CHARMM36 systems, the preparation included energy minimization, followed by 500 ps each of NVT and NPT equilibration. For the Amber systems, two NPT equilibration steps followed the initial NVT equilibration: a 100 ps equilibration with a 1 fs timestep and a subsequent 1 ns equilibration with a 2 fs timestep. These simulations were performed at 300 K using Langevin dynamics. Long-range electrostatics were computed using the particle-mesh Ewald (PME) method, with a grid spacing of 1.2 Å^76^, and a nonbonded cutoff of 9 Å was applied.

Production simulations were conducted for both force fields with two replicas per system. The total simulated times were 2.67 μs and 2.56 μs for CHARMM36 systems, and 3.25 μs and 3.06 μs for Amber systems (Table 1). The solvent-accessible surface area (SASA) of the TF protein was computed for all simulations using the gmx sasa program.

**Table 1.** Details of the molecular dynamics simulations.

| System | # water molecules | # atoms | Replicas | Simulation time (per replica) |
| --- | --- | --- | --- | --- |
| TF (monomer) (CHARMM) | 33737 | 108229 | 2 | 2680 ns |
| TF (monomer) (AMBER) | 34373 | 110205 | 2 | 3250 ns |
| TF + RNC | 108382 | 337347 | 3 | 1000 ns |
| TF + L17 | 26015 | 86167 | 1 | 800 ns |
| TF + L23 | 36111 | 116098 | 1 | 800 ns |

For all the simulations VMD software^77^ was used for analysis of the trajectories and production of some of the figures. PyMol software (Pymol) was used for figure production.

#### b. Trigger Factor Binding to the Ribosome Nascent Chain Complex

The interaction between the RNC and TF was predicted using AlphaFold Multimer^48,78^ and the model with the best score was prepared for simulation. Briefly, hydrogen atoms were added to the complex using the VMD software^76^. The system was then solvated resulting in a cubic box of dimensions described in Table X1. K^+^ and Cl^-^ ions were added using the system resulting in an ionic concentration of 0.15 M KCl. The preparation included energy minimization, followed by NVT and NPT equilibration, first at 1 fs timestep for 250 ps each, and another 250 ps for the NPT simulation at 2 fs. Three individual replicas were run for 500 ns.

These simulations were performed with NAMD3.0.1^79^. The CHARMM36 force field^70^ was used to model the protein and ions, and the TIP3P model^71^ was chosen for the water. The PME method^76^ was used for the treatment of electrostatic interactions. Electrostatic and van der Waals forces were calculated in every time step with a 12 Å cutoff distance. A switching distance of 10 Å was chosen to smoothly truncate the non-bonded interactions. The Nose-Hoover-Langevin piston method was employed to control the pressure with a 50 fs period, 25 fs damping constant, and a desired value of 1 atmosphere. These simulations were performed at 298 K using Langevin dynamics.

Distances between the ribosome nascent chain and C-terminal of TF were measured by computing the minimum distance between pairs of heavy atoms in the interaction interface throughout the trajectories of the three replicas. This interaction interface was defined as residues 286 to 355 for the ribosome nascent chain and residues 300 to 392 for the TF. The same approach was used to compute minimum distances between RNC and residues 320 and 377 of TF.

#### c. Ribosome-Trigger Factor docking and MD simulations

Rigid body docking between ribosomal proteins (L17 and L23) and theTF monomer was performed using Piper^80^. The ribosomal proteins were extracted from the whole ribosome structure (PDB 7K00) and docked in isolation with the TF monomer (PDB 1W26). Some poses generated through docking were deemed unrealistic, as they positioned TF in regions inaccessible within the complete ribosome structure. Therefore, for each ribosomal protein, physically implausible poses were filtered from the top 15 poses generated.

MD simulations of the remaining poses were carried out using GROMACS 2020^74^ with the CHARMM36 force field^70^ (Table 1). The systems were solvated with TIP3P water molecules^71^, and neutralized to an ionic concentration of 0.15 M. The MD protocol consisted of four stages: energy minimization, NVT equilibration, NPT equilibration, and production.

During equilibration, a 500 ps NVT simulation at 300 K was performed using Langevin dynamics with a 2 fs timestep. Long-range electrostatics were treated using the PME method^76^, with a grid spacing of 1.2 Å, and a nonbonded cutoff of 12 Å. Positional restraints were applied to the protein alpha-carbons during this step. Following NVT equilibration, a 500 ps NPT equilibration was conducted at 1 bar using the Parrinello-Rahman barostat^81^. Production runs of 100 ns were performed for each system starting from the equilibrated structures.

The simulations were analyzed to characterize the binding interfaces, defined as residues within 6 Å of the opposing protein. Metrics included the percentage of preserved contacts relative to the final frame, the distance between the centers of mass of both interfaces, and the RMSD of the ribosomal and TF interfaces. These metrics were used to assess stability and identify unbinding events or instability in docking-predicted poses. Specifically, a significant increase in the distance between the centers of mass indicated unbinding, while slight decreases reflected interface rearrangements resulting in more stable conformations not captured during docking. Poses with stable center-of-mass distances but low contact preservation or high RMSDs were also considered unstable.

The results of these initial simulations served as an additional filter to select the most stable docking poses. Stable poses were subjected to extended simulations of ∼800 ns each to validate their stability further. The metrics from these extended simulations were used to rank the remaining poses, and the most stable one for each ribosomal protein was used as the final model for analysis.

## Acknowledgements

We gratefully acknowledge the Basque Government for their funding and support through the grants (POS_2021_1_0017) and (IT-1707-22), as well as the technical and human support provided by the DIPC Supercomputing Center. We also thank the Spanish Ministry of Science and Innovation for the project (PID2022-139230NB-I00). This project has received funding from the European Research Council (ERC) under the European Union’s Horizon 2020 research and innovation programme (grant agreement No 772257).

---

**Supplementary Figure 1.**
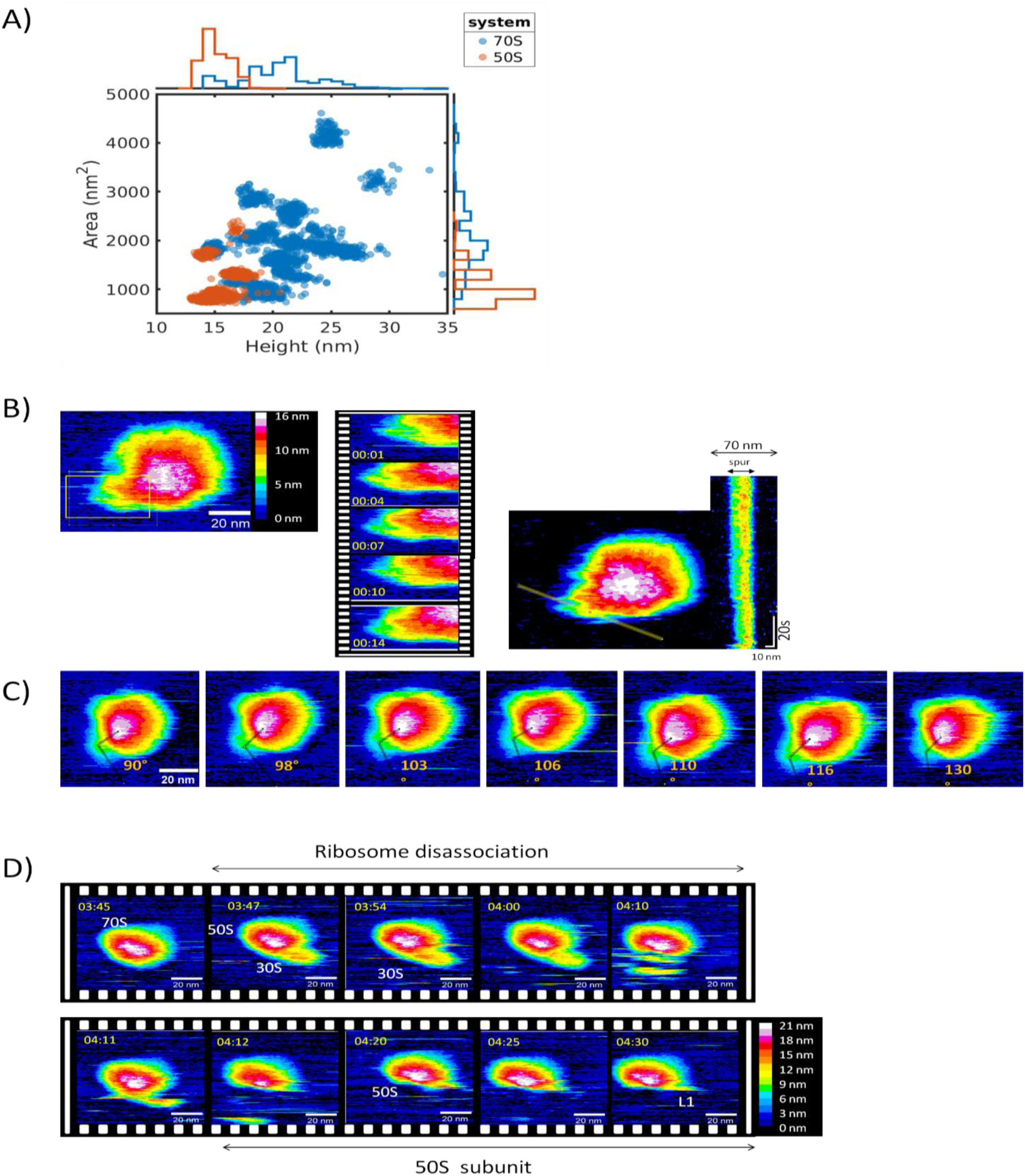
Distribution and Structural Dynamics of 70 S Ribosome. **(A)** Plot showing the relationship between ribosome height (X-axis, in nm) and area (Y-axis, in nm²) as measured by HS-AFM. The histograms above and to the right of the plot display the height and area distributions, respectively. Blue corresponds to the 70S ribosome, and orange represents the 50S subunit. **(B)** Left: Structural dynamics of the spur on the 30S subunit, with sequential frames highlighting spur movement. Right: Kymograph of the yellow line (spur) drawn across the HS-AFM image frames. **(C)** Structural dynamics of the ribosomal P-stalk base of the 50S subunit. The angles between the proximal and distal regions of the stalk are indicated by the yellow line (θ) in the four images. **(D)** Sequential frames illustrating the dissociation of the 70S ribosome into its 50S and 30S subunits.

**Supplementary Figure 2.**
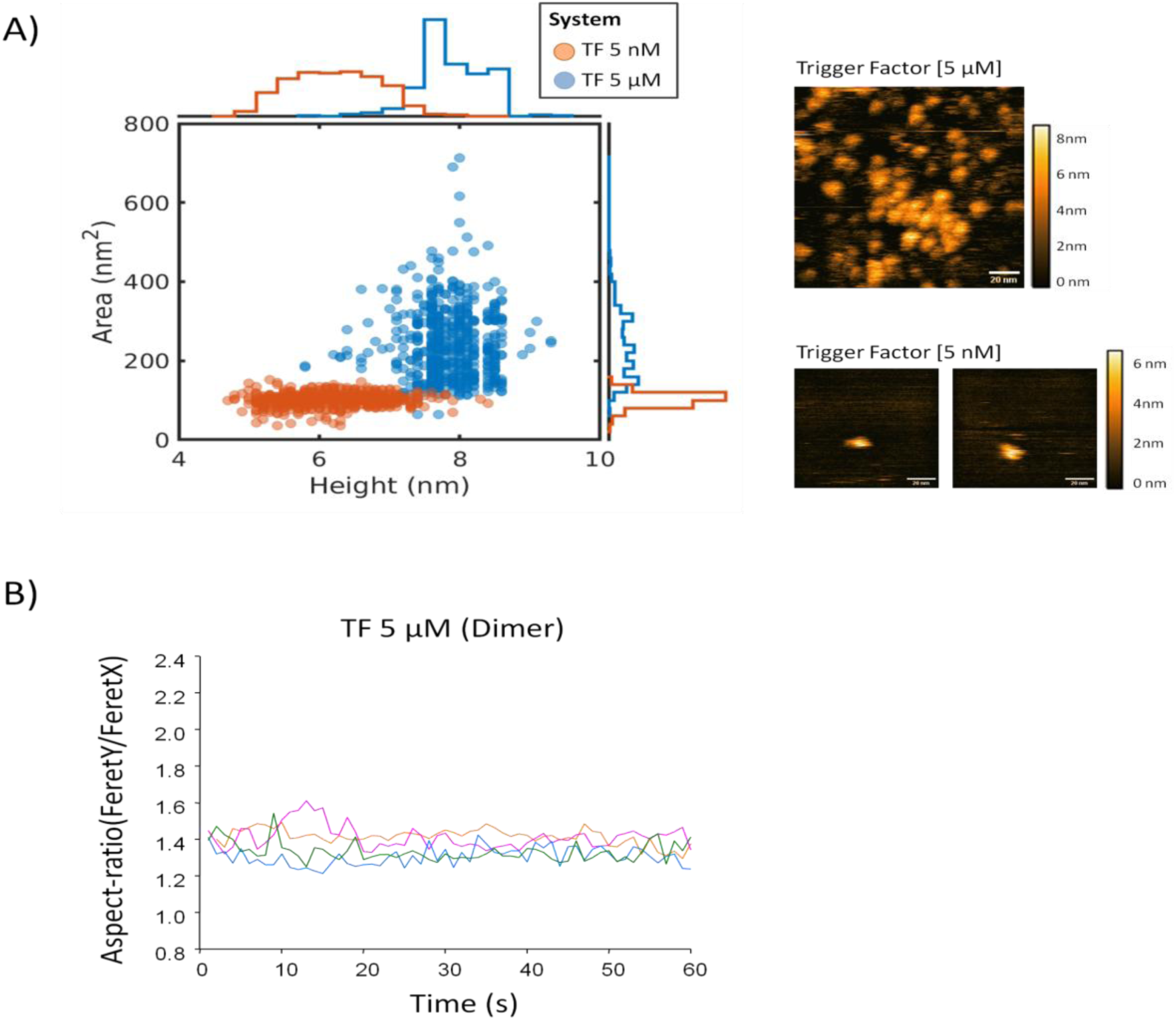
TF Monomer and Dimer Distribution and Aspect Ratio Variation. **(A)** Plot showing the relationship between the height and area of Trigger Factor (TF) as measured by HS-AFM. The histograms above and to the right of the plot display the height and area distributions, respectively. Orange represents data from TF at 5 nM, and blue of TF at 5 µM. Top right: A frame showing TF at 5 µM. Bottom: Two frames of TF at 5 nM in different conformational states. **(B)** Time-dependent variation of the aspect ratio of TF dimers during HS-AFM visualization. Data from different experiments are shown in distinct colors. Measurements were performed at a 5 µM concentration in physiological buffer (25 mM Tris, pH 7.5, 120 mM KCl, 5 mM NaCl, 14 mM MgAc).

**Supplementary Figure 3.**
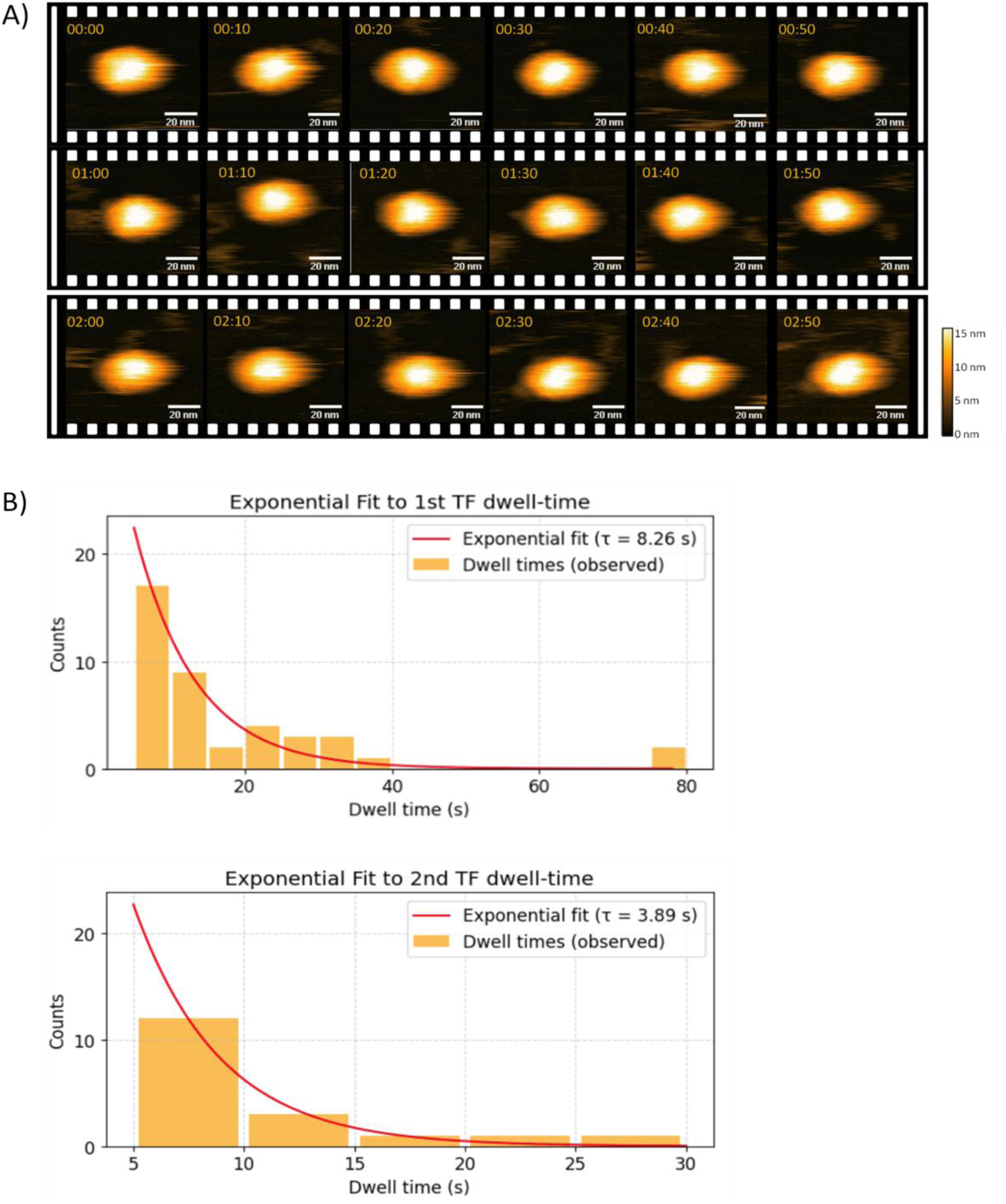
Temporal dynamics of TF interactions with non-translating ribosomes. **(A)** Sequential frames from HS-AFM showing the 70S ribosome in the absence of translation, in the presence of 1 µM TF, without detectable binding of TF to the ribosome. **(B)** Dwell time of TF interaction with the RNC complex and exponential distribution fit. The top histogram shows the observed dwell times of the first TF molecule (in orange, as counts), with the red curve representing the exponential fit (τ = 8.3 ± 1.1 seconds). The bottom histogram corresponds to the dwell time of a second TF molecule interacting simultaneously with the ribosome. The red curve indicates the exponential fit, yielding a shorter time constant of τ = 3.9 ± 0.4 seconds.

**Supplementary Figure 4.**
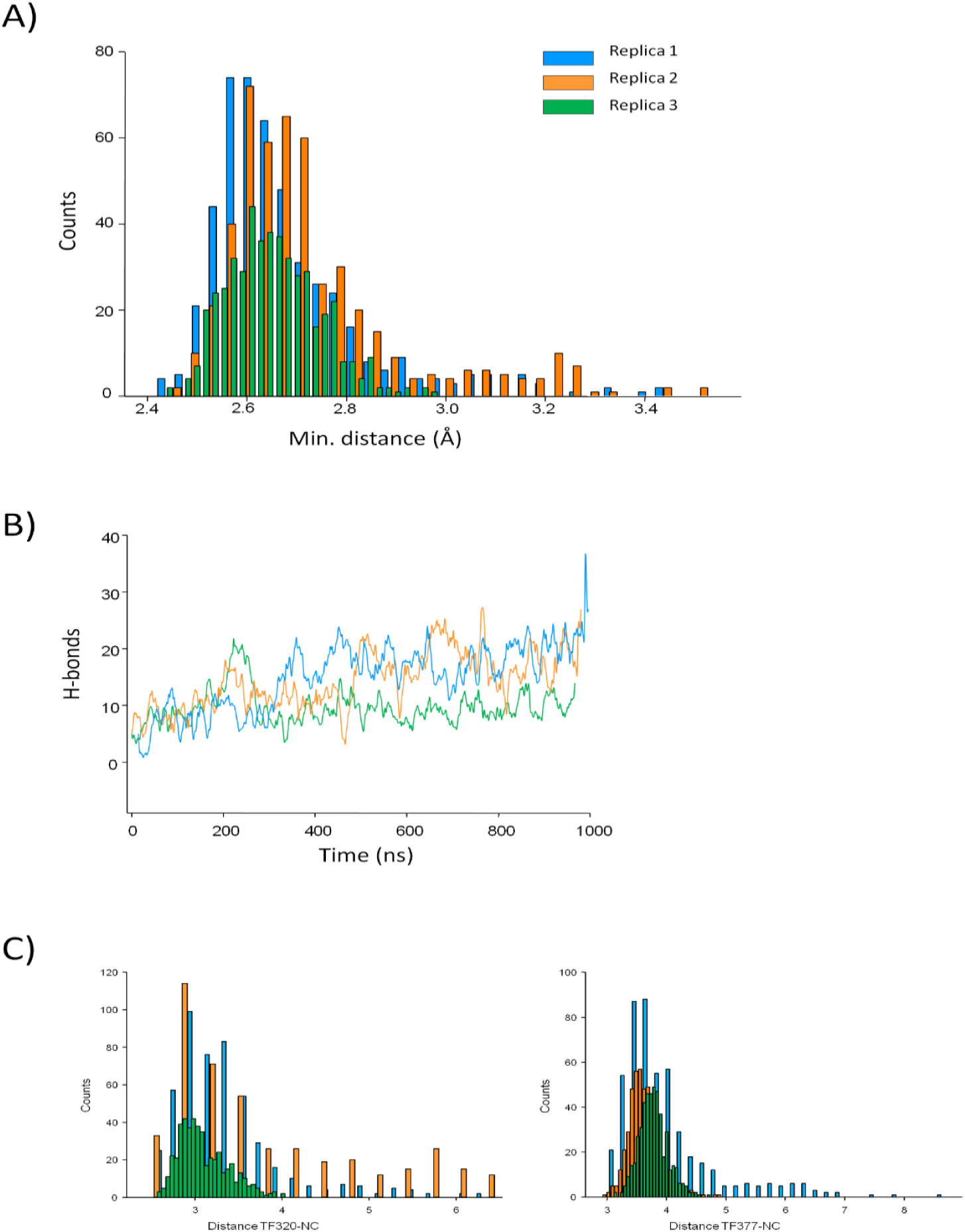
Temporal dynamics of the interaction between TF and the nascent chain, based on all-atom molecular dynamics simulations. **(A)** Histogram showing the minimum distance between the C-terminal of TF (residues 300–392) and the hA-TW-B region of the nascent chain. Data from different replicates are shown in distinct colors. **(B)** The number of hydrogen bonds between the two chains is shown across three replicates, with each replicate represented by a different color. The curves represent a moving average of the raw data. Data from different replicates are shown in distinct colors. **(C)** Histogram showing the minimum distance between the TF320 (left) and TF377 (right) and the hA-TW-B region of the nascent chain. Data from different replicates are shown in distinct colors.

**Supplementary Figure 5.**
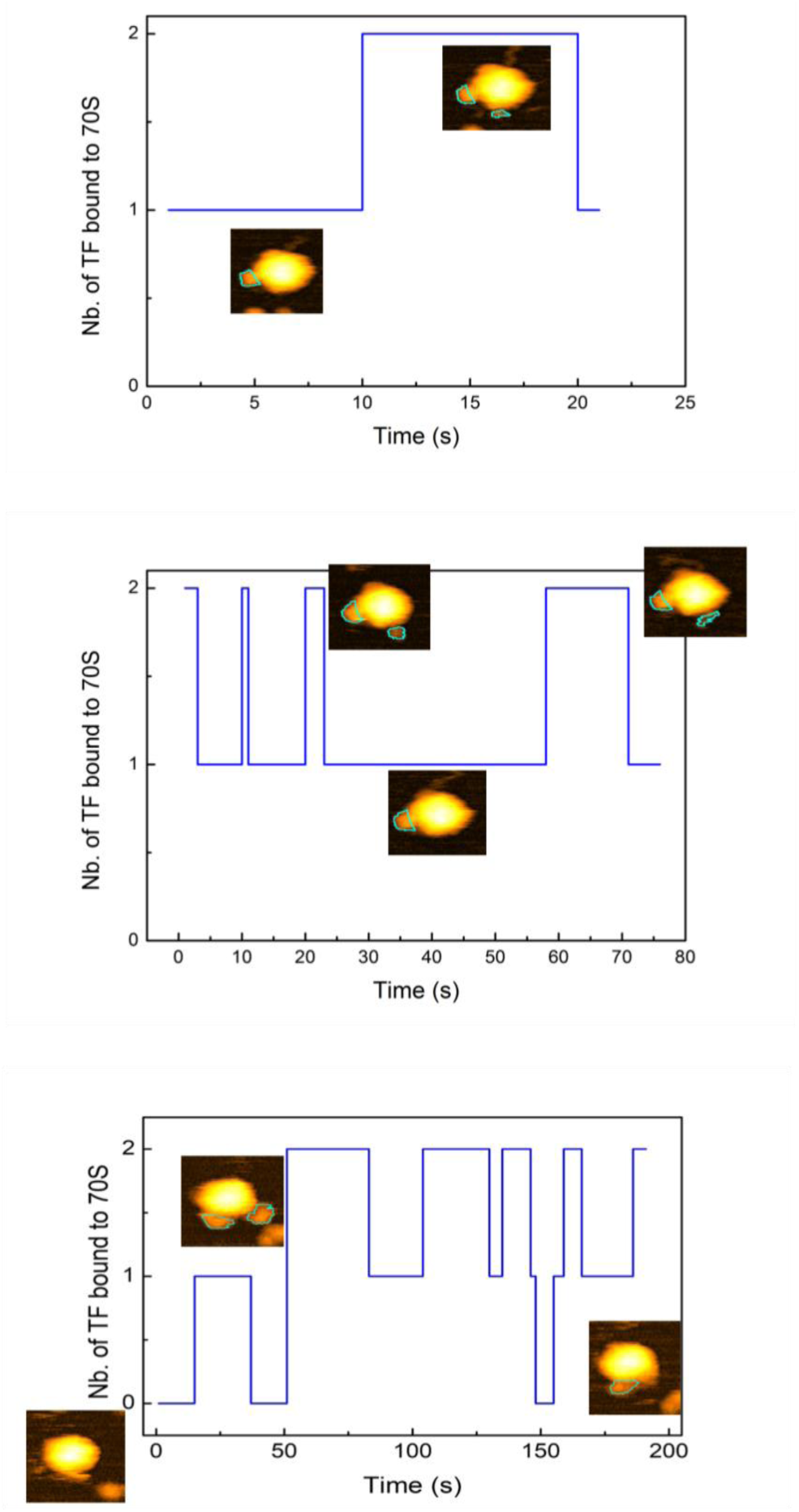
Simultaneous Binding of Multiple TF Molecules to the Ribosomes. Graph showing the number of TF molecules bound over time, highlighting the frequency and duration of binding events. Data from three distinct video recordings, each featuring the RNC in different orientations, are presented. The encircled particles in the topographs show the detected binding events.

